# CTCF association with episomal HPV16 genomes regulates viral oncogene transcription and splicing

**DOI:** 10.1101/2021.02.18.431881

**Authors:** Ian J Groves, George Tang, Ieisha Pentland, Joanna L Parish, Nicholas Coleman

## Abstract

Human papillomavirus 16 (HPV16) is a high-risk alphapapillomavirus that is associated with cancers of mucosal epithelia. The virus genome exists in cells as an episome but can integrate and overexpress the *E6* and *E7* viral oncogenes. In related high-risk family members HPV18 and HPV31, host proteins including CTCF, an insulator, and SMC1A, a component of the cohesion complex, are known to interact with the viral genome and alter transcriptional activity, splicing patterns and episome amplification. However, the roles of these two proteins during HPV16 infection has not yet been fully examined. Here, we show during differentiation of the episomal HPV16-containing W12 cell line that CTCF association increases with the virus genome at the known *E2* binding site, whilst additional CTCF binding now occurs at the putative *L2* binding site, with SMC1A association occurring unchanged here. While expression of virus late transcripts (*E4^L1, L2, L1*) is stimulated, early transcript levels decrease by 48 hours, with the exception of the E6*IV spliced transcript. Conversely, in undifferentiated, monolayer W12 cells, CTCF knockdown increases the level of all early transcripts, whereas E6*IV level increases. Additionally, CTCF ablation as well as SMC1A knockdown results in decreases to HPV16 genome copy number. Taken together, this supports the model that while CTCF and SMC1A have a role in HPV16 genome maintenance, CTCF plays a greater part in regulating HPV16 oncogene splicing and expression during the natural lifecycle of the virus, and may be involved in a reduced risk of cancer development during episomal HPV16 infections.

## Introduction

Human papillomaviruses (HPVs) are small, non-enveloped DNA viruses that infect mucosal and cutaneous epithelia[1]. High-risk human papillomavirus (HRHPV) infections are associated with over 5% of all human cancers[2], including cervical, vaginal, vulvar, penile, anal, and head and neck malignancies[3, 4]. Importantly, alone HPV types 16 (HPV16) and 18 (HPV18) are responsible for around 70% of cervical cancers[5], which routinely develop from squamous intraepithelial lesions (SILs)[6] into squamous cell carcinomas (SCCs)[3].

During productive virus infection, HPV enters the basal layer of stratified epithelia and expres ses early genes, including oncogenes *E6* and *E7*, from the viral early promoter[3]. However, as cells differentiate and move to more superficial layers, E6 and E7 protein levels decrease and late genes are expressed through initiation of transcription from the late promoter[7]. Dysregulation of this control, and virus oncogene expression, is an essential event in HPV-associated carcinogenesis that regularly occurs due to integration of the virus genome into that of the host[8]. However, ∼15% of cervical SCCs are known to carry episomal HPV16 genomes from which deregulated oncogene expression still occurs[9].

Gene expression from the HPV episomal genome during the virus life cycle is regulated by both viral and multiple host proteins during differentiation of infected cells[10]. Alternative splicing and polyadenylation is essential for HRHPVs, as multiple genes are transcribed from each promoter as polycistronic transcripts[11], which are then spliced into separate mature transcripts to encode individual viral proteins[12]. A major contrast between high- and low-risk alphapapillomaviruses is that HRHPVs encode E6 star (E6*) products that use different splice donor and acceptor sites in order to synthesise varying early gene products, including E6 and E7 proteins[6]. One host protein that is known to regulate alternative splicing in HRHPVs is CCCTC-binding factor (CTCF)[13]. CTCF is a ubiquitously expressed host protein that binds to DNA at sequence-specific motifs through 11 zinc fingers, a common motif in DNA-binding proteins[14], and control of binding to host target sites is mediated through DNA methylation of cytosines on CpG dinucleotides[12]. CTCF has been shown to have roles both in the control of transcriptional regulation and DNA replication of many virus families, including adeno-, herpes-, polyoma-(Groves, unpublished data) and papillomaviruses[12].

CTCF has previously been shown to bind to genomes of HRHPVs through conserved CTCF binding sites in the open reading frame (ORF) of *E2* and also the *L2* gene of HPV31[13, 15]. Studies of episomal HPV18 have shown that binding of CTCF to the *E2* ORF regulates viral gene expression during epithelial differentiation, with disruption of CTCF binding causing an increase in the level of unspliced early transcripts[13]. Loss of CTCF binding also caused a large decrease in the level of a HPV18 spliced transcript (E6*II, 233^3434), which encodes the E6 and E5 proteins[13]. Further work has shown that CTCF-mediated control of HPV18 gene expression occurs through interaction with the host Yin Yang 1 (YY1) protein at the virus long control region (LCR) such that virus chromatin is looped, with the virus enhancer epigenetically repressed through polycomb repressor complex (PRC) recruitment[16]. Cellular differentiation results in decreasing YY1 expression, disruption of HPV18 genome looping and epigenetic repression leading to increased viral gene expression[16].

A further host protein that has been investigated for its role in replication of HPV31 episomal genomes is SMC1A, a cohesin protein[15]. SMC1A was shown to interact with the late region of HPV31, through CTCF, to alter genome stability, with the authors suggesting that SMC1A has a role in aligning viral genomes in G2 phase of mitosis towards facilitating genome duplication[15]. However, the association of SMC1A or the effects of altered CTCF interaction with HPV16 genomes remains unstudied. Here, we show that during methylcellulose-induced differentiation of HPV16 episome-containing W12 cells, CTCF enrichment at the virus genome increases concomitant with modulation of viral gene expression and induction of late gene expression. SMC1A interaction occurs at the *E2* CTCFbs but remains unchanged during differentiation. Additionally, CTCF ablation in monolayer, undifferentiated W12 cells causes an increase in early gene transcript levels but, remarkably, a decrease to the HPV16 E6*IV spliced transcript, confirming the role of CTCF in transcriptional and splicing control during the episomal HPV16 life cycle.

## Materials and Methods

### Cell culture and differentiation

The W12 cell li ne, derived from a CIN 1 lesion of the cervix, has the properties of basal epithelial cells and typically carries ∼100-200 HPV16 genomes per cell[17, 18]. The generation of these cells has been described previously[19]. The original episomal population of W12 cells is polyclonal and is described elsewhere as the ‘parental’ (Par1) cell line[9]. Monolayer cell passage was performed using protocols and reagents that have been described elsewhere[20]. For methylcellulose differentiation, cells were seeded at ∼1×10^6^-1×10^7^ cells/ml of 1.6% methylcellulose in culture medium and incubated up to 48h. For harvest, cells were washed via centrifugation with excess 1xPBS at least three times to remove all traces of methylcellulose medium.

### Gene depletion

Each target gene was depleted using human ON-TARGETplus SMARTpool siRNAs (GE Dharmacon): non-targeting control (NTC, D-001810-10); CTCF (L-020165-00); SMC1A (L-006833-00). All siRNAs were used at a final concentration of 10 nM (including dual ablation of CTCF/SMC1A), with cells being transfected at 20 –30% confluence using Lipofectamine RNAiMAX (Invitrogen) as described[21-23] for 48 hr before analysis of target knockdown and effects on HPV16 transcription.

### Quantification of proteins, transcripts and virus genome copy number

Quantitative immunoblotting was carried out as described previously[9, 24, 25] using the antibodies listed in Table 1. Protein concentrations for CTCF and SMC1A were compared with those of the β-actin loading control using ImageJ software analysis. Levels of host and virus transcripts were measured using SYBRGreen quantitative reverse transcription–PCR (qRT–PCR) as described[25]. Primers and conditions are given in Table 2 or published previously[25, 26]. Relative transcript levels per cell were determined using the Pfaffl equation[27], normalised to the mean of four housekeeping genes (ACTB, GAPDH, RPL13A and YWHAZ)[28], then referenced to control samples. Relative transcript levels per HPV16 template were determined via quantification to the HPV16 genome copy number per cell. For HPV16 genome copy number analysis, DNA extraction was carried out as previously described[29] before qPCR was used to determine mean *E6* (111F to 223R), *E7* (649F to 765R), E2 5’ (2695F to 2757R) and E2 3’ (3936F to 4025R) DNA level in comparison to mean host *GAPDHprom* and *MyoGprom* regions, with analysis as previously described[26].

**Table 1.** Summary of protein targets for ChIP and Immunoblot analysis.

| Protein target | Supplier | Cat. number | Protocol | Volume / dilution |
| --- | --- | --- | --- | --- |
| CTCF | Millipore | 17-10044 | IB | 1 µg/ml |
|  | Active Motif | 61311 | ChIP | 8 µl |
| SMC1A | Abcam | ab9262 | IB | 1 µg/ml |
|  | Active Motif | 61067 | ChIP | 4 µl |
| β-Actin | Abcam | ab6276 | IB | 31 ng/µl |
| H3K4me3 | Active Motif | 39915 | ChIP | 5 µl |
| IgG Mouse | Millipore | 12-371 | ChIP | Variable |
| IgG Rabbit | Dako | X0902 | ChIP | Variable |

**Table 2.**
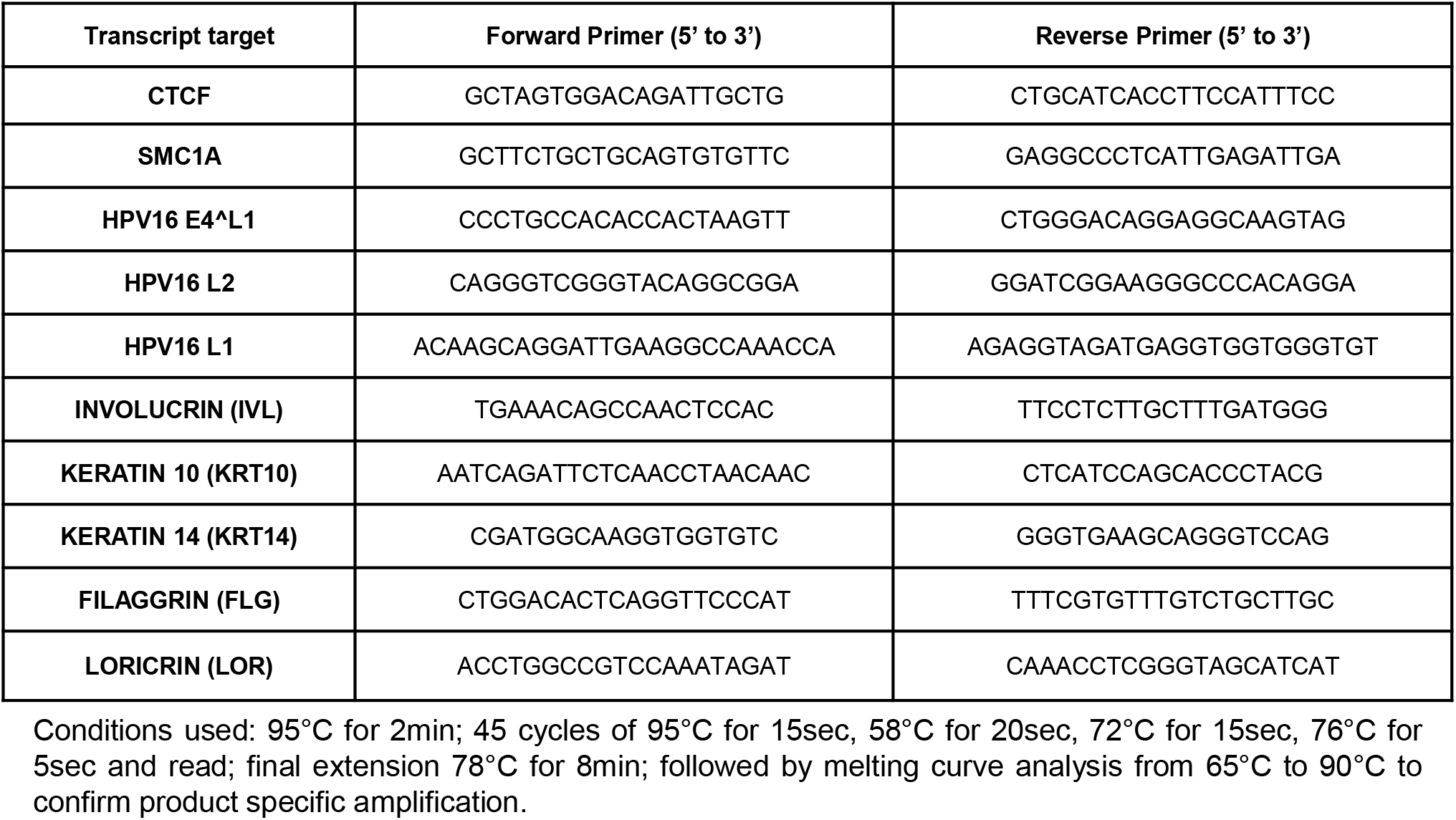
Primers and conditions for qRT-PCR.

### Chromatin immunoprecipitation

Chromatin immunoprecipitation (ChIP) was performed as described[25, 26] using ChIP-IT Express kits (Active Motif) and ChIP-validated primary antibodies for CTCF, SMC1A and the active transcription histone post-translational modification H3K4me3 with appropriate serum/IgG negative controls (Table 1). Enrichment of DNA across the full length HPV16 genome was then assessed using qPCR, primers and conditions in comparison to negative controls as described previously[26].

## Results

### Differentiation of W12 cells causes modulation of episomal HPV16 spliced transcript levels

To investigate the changes to early and late transcripts from the HPV16 genomes in W12 cells during differentiation, we first assessed the initial transcript levels in monolayer, undifferentiated cells (Fig. 1). Using a range of primer pairs that would detect all transcripts encoding the E6 and E7 proteins, as well as the alternatively spliced forms, E6*I, E6*II, E6*III, E6*IV, E6*X, and both 5’ and 3’ ends of E2 transcripts[25, 30], we found highest expression of total E6 transcripts (E6all), E7, E1*I and E6*X (Fig. 1C). There was also a high level of E2 3’ transcript, likely due to all early transcripts terminating at the early polyadenylation site just downstream of the 3’ end of the *E2* gene, and consistent with previous W12 episomal RNA-seq data[31]. All other transcripts were minimally expressed, including those encoding late proteins (*E4^L1, L2* and *L1*).

**Figure 1.**
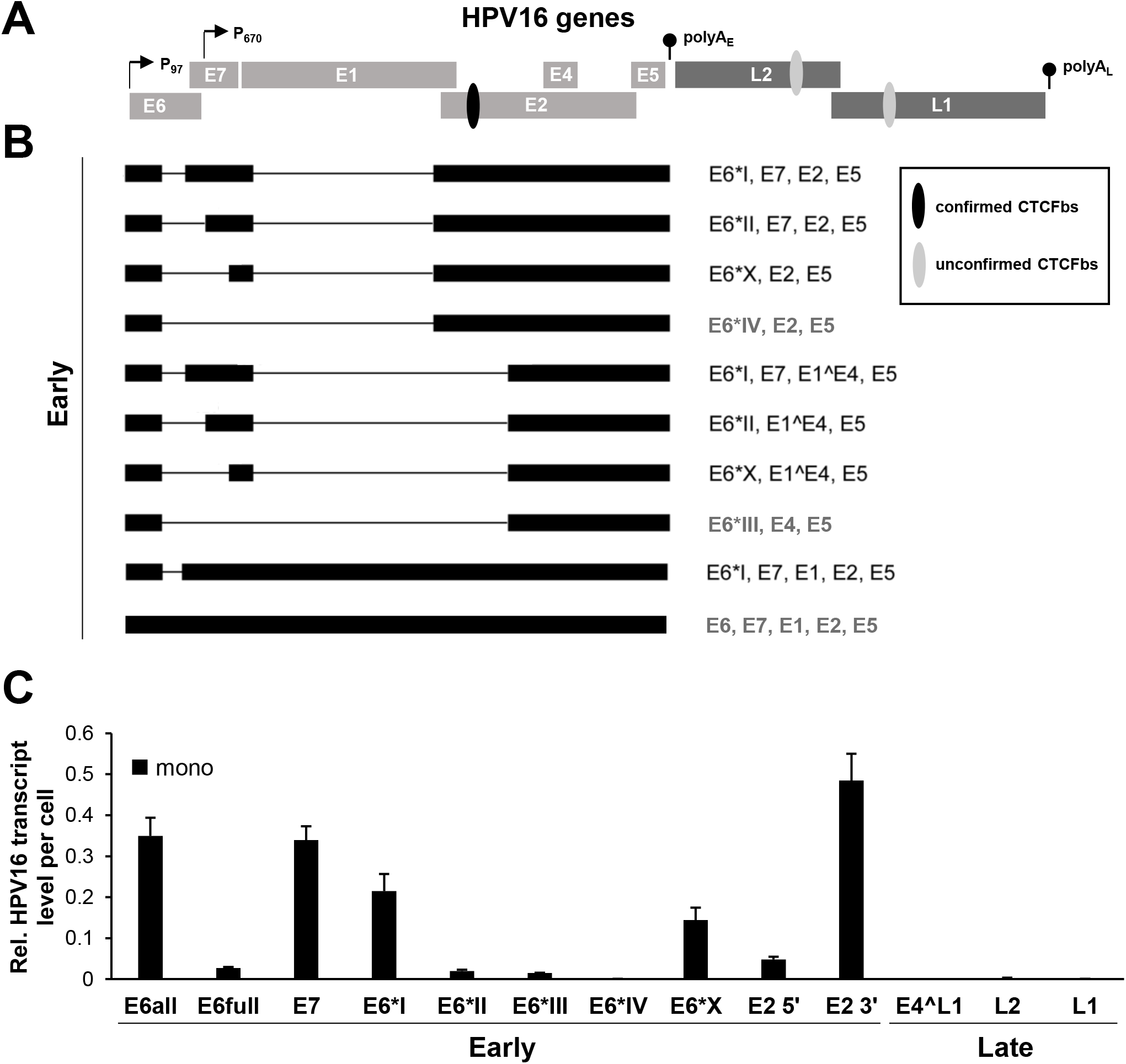
HPV16 transcripts in undifferentiated monolayer W12 episomal cells. Schematic representation of (A) the HPV16 genes with confirmed CTCF binding sites (black oval) and unconfirmed CTCF binding sites (grey ovals) and (B) the major HPV16 transcripts that terminate at the early polyadenylation site in W12 cells analysed in this study (adapted from http://pave.niaid.nih.gov). (C) Level of early and late (E4^L1, L2, L1) HPV16 transcripts found in undifferentiated monolayer W12 cells, relative to host housekeeping genes. Bars = mean+s.e.m. of n=3. (P = promoter; polyA_E_ = early polyadenylation site.)

We next differentiated W12 cells using methylcellulose culture conditions for 24 and 48 hours before analysing HPV16 transcript levels (Fig. 2). Differentiation of cells was confirmed by analysis of transcript changes to cellular marker genes, with late differentiation transcripts Involucrin, Filaggrin and Loricrin all increasing, whilst early transcripts Keratin 10 and 14 decreased (Fig. 2B) in comparison to levels found in monolayer cells (Fig. 2A). In addition, analysis of HPV16 genome copy number showed increases by 24 (1.24-fold) and 48 hours (2.13-fold) post-methylcellulose treatment versus monolayer cells (Fig. 2C). Differentiation resulted in an initial increase to all transcripts (early and late) per cell by 24 hours post-treatment, which waned by 48 hours in most cases, causing an overall decrease in E6*III level (Fig. 2D). However, whilst both E2 transcripts followed an increasing trend in line with the induction of late transcripts, E4^L1, L2 and L1, surprisingly E6*IV increased 62.2-fold. By taking account of increasing virus genome copy number, all early transcripts could be seen to decrease below monolayer levels, excluding the E2 transcripts that declined slightly from their 24 hour peak, and E6*IV whose rise was clearly maintained (Fig 2E).

**Figure 2.**
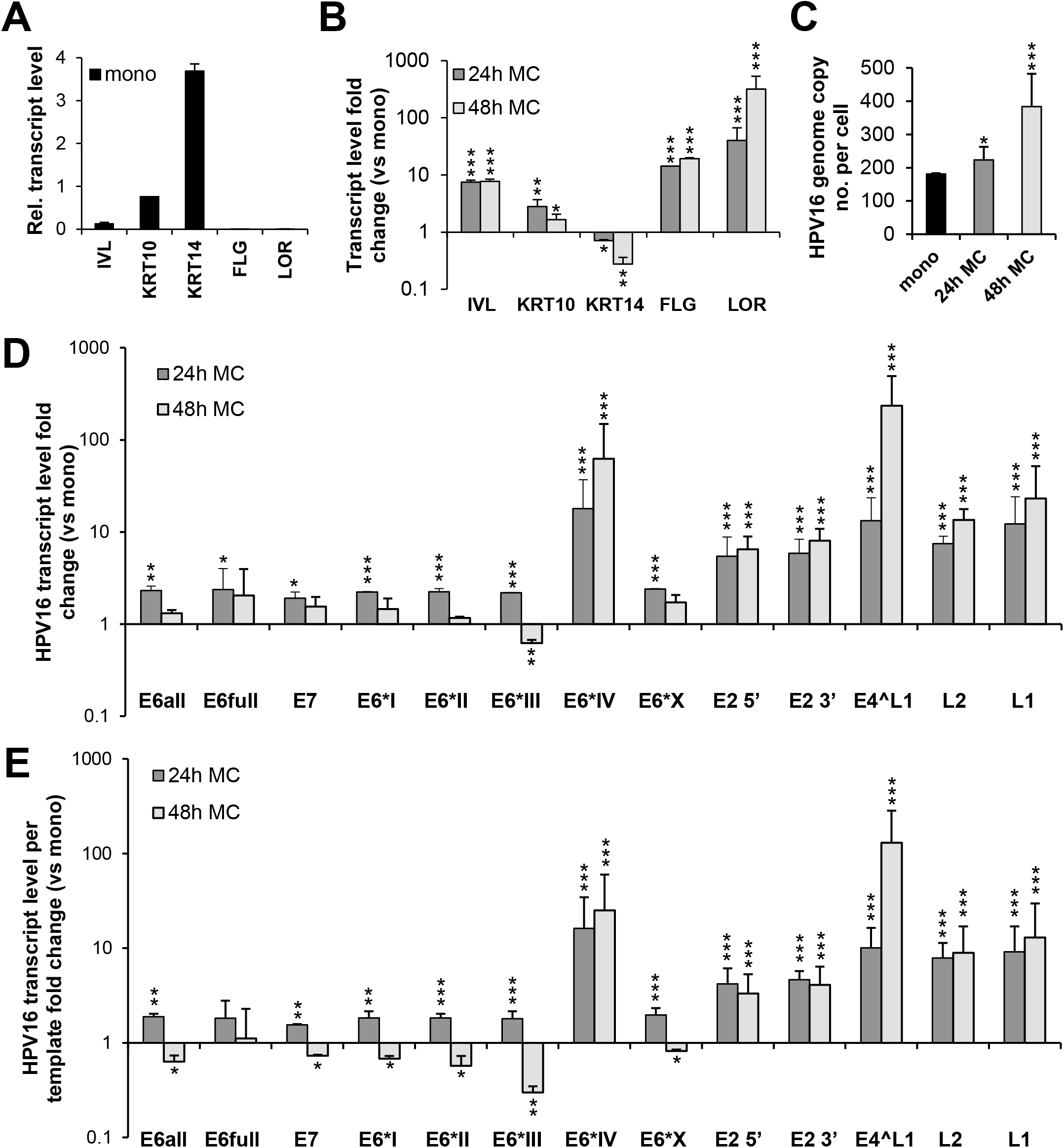
Modulation of HPV16 transcript level in W12 episomal cells during differentiation. (A) Relative level of epithelial differentiation marker transcript level in undifferentiated monolayer W12 episomal cells (mono, black bars) and fold-changes of level with methylcellulose differentiation (24h MC, dark grey bars; 48h MC, light grey bars) with (C) level of HPV16 genome copy in the associated cells. (D) Fold-change level of early and late HPV16 transcripts per cell with methylcellulose differentiation (24h MC, dark grey bars; 48h MC, light grey bars) (relative to monolayer cells, Fig. 1D) with (E) HPV16 copy number-accounted transcript fold-change levels. Bars = mean+s.e.m. of n=3. Student’s T-test vs monolayer data: *p<0.05, **p<0.01, ***p<0.001. (IVL, Involucrin; KRT10, Keratin 10; KRT14, Keratin 14; FLG, Filaggrin; LOR, Loricrin.)

### Differentiation-mediated modulation of episomal HPV16 transcription is associated with increased association of CTCF at the virus genome

To interrogate the association of host proteins known to interact and regulate transcription from high-risk HPV episomal genomes, we next carried out chromatin immunoprecipitation (ChIP) analysis on monolayer and methylcellulose treated episomal W12 cells (Fig. 3). Cohesin protein SMC1A was found at the *L2* CTCFbs binding site before and after differentiation, with only a slight enrichment increase across the *L1* CTCF binding site and *E1* gene by 48h (Fig. 3B). However, CTCF association increased significantly at, and between, the *E2* CTCFbs and *L2* CTCFbs after 24 and 48 hours methylcellulose treatment, with a weak (but significant) concurrent increase over the HPV16 LCR (Fig. 3A). In contrast, a histone post-translational modification (PTM) of active transcription, H3K4me3, was enriched above levels seen in monolayer, undifferentiated W12 cells only after 48 hours of methylcellulose treatment across the late promoter and *E1* gene (Fig. 3C).

**Figure 3.**
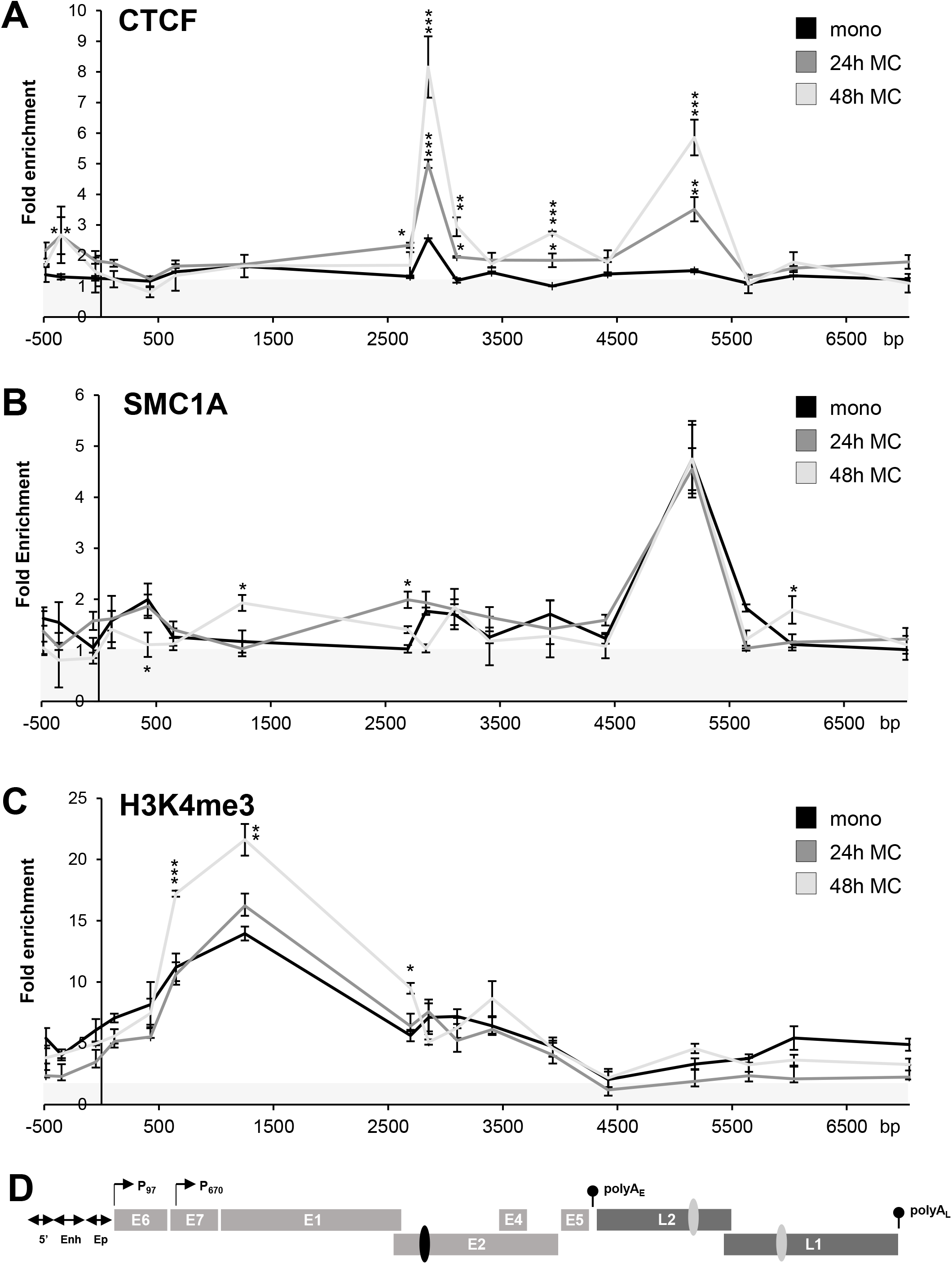
Differentiation of W12 episomal cells causes increases in CTCF association with HPV16 genomes. Chromatin immunoprecipitation analysis of (A) CTCF, (B) cohesion SMC1 and (C) active transcriptional mark H3K4me3 across (D) the episomal HPV16 genome from undifferentiated monolayer (mono, black line) and methylcellulose differentiated (24h MC, dark grey line; 48h MC, light grey line) W12 episomal cells. In each graph, the y-axis shows the relative levels of enrichment above IgG controls (<1, grey area). Data points = mean±s.e.m. of n=3. Student’s T-test vs monolayer data: *p<0.05, **p<0.01, ***p<0.001.

### CTCF protein ablation results in HPV16 genome decreases but early transcript increases, excluding spliced E6*IV

As modulation of HPV16 transcript levels during cellular differentiation had coincided with increasing association of CTCF at the HPV16 genome, we next assessed whether CTCF protein ablation in monolayer, undifferentiated episomal W12 cells would affect HPV16 transcript profiles (Fig. 4). Ablation of both CTCF and SMC1A individually and together (CTCF/SMC1A), using small interfering RNA (siRNA) pools, was confirmed by transcript (Fig. 4A, B) and protein (Fig. 4C, D) analysis. Next, HPV16 genome copy number was determined after 48 hours siRNA treatment, and decreases seen with both individual (CTCF, 18% down; SMC1A, 42% down) and dual ablation (CTCF/SMC1A, 31% down) treatments in comparison to non-targeting control (NTC) siRNAs (Fig. 4E). Loss of association of CTCF with the HPV16 genome was confirmed by ChIP analysis, where enrichment at the *E2* CTCFbs in monolayer W12 cells was lost with CTCF-specific siRNA treatment in comparison to NTC siRNAs (Fig. 5A). Unfortunately, ablation of SMC1A at the HPV16 genome could not be confirmed due to failure of ChIP analysis with such a reduced number of virus genomes. However, transcript analysis could be conducted with the sample material from all treatments (Fig. 6). Although only moderate, a significant increase of all HPV16 early transcripts apart from E7 and E2 5’, occurred with CTCF ablation but, conversely, a decrease to E6*IV (40% down), with no change seen to late transcripts (Fig. 6A). SMC1A ablation alone caused no significant changes to any HPV16 transcripts, although E6*IV level was again decreased (Fig. 6B). Dual ablation of CTCF/SMC1A (Fig. 6C) gave no change to HPV16 transcript levels beyond that of CTCF alone, however E6*IV was unchanged from control levels. When taking account of HPV16 copy number decreases, all treatments now caused early transcript level increases, though changes to late gene expression remained non-significant (Fig. 6B, D, F). Uniquely, E6*IV level decrease was seen only with CTCF ablation (Fig. 6B).

**Figure 4.**
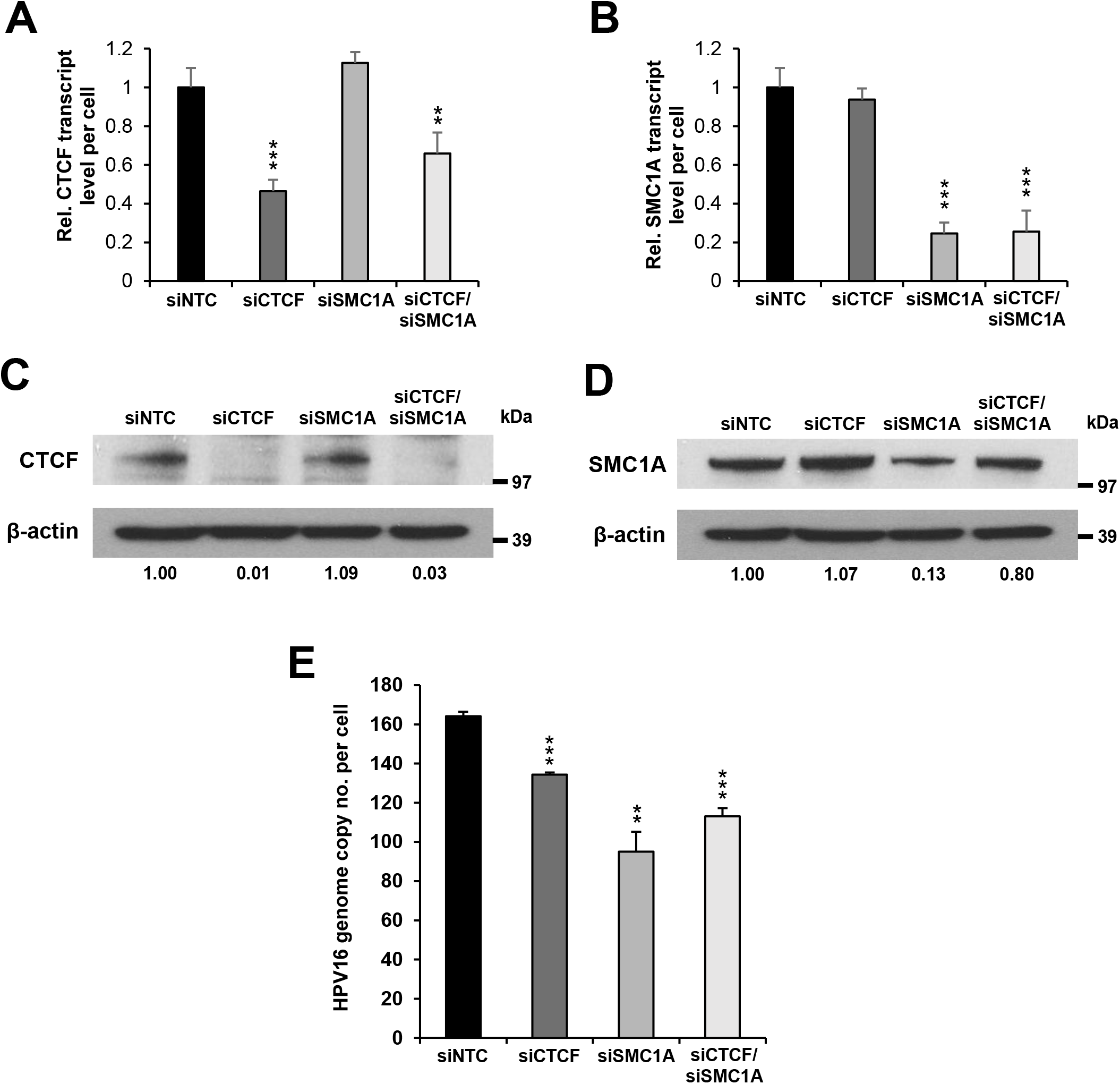
CTCF and SMC1A knockdown causes HPV16 genome copy number loss in undifferentiated monolayer W12 episomal cells. Monolayer W12 episomal cells were treated with non-targeting control (NTC, black bars) or siRNAs specific to CTCF (dark grey bars), SMC1A (medium grey bars) and CTCF/SMC1A (light grey bars) with depletion level established of (A, B) transcripts and (C, D) protein (NTC set to 1), and (E) level of HPV16 genome copy per cell determined in the associated cells. Immunoblots are representative of n=3, with ImageJ quantified values vs β-actin shown below each panel. Bars = mean+s.e.m. of n=3. Student’s T-test vs monolayer data: **p<0.01, ***p<0.001.

**Figure 5.**
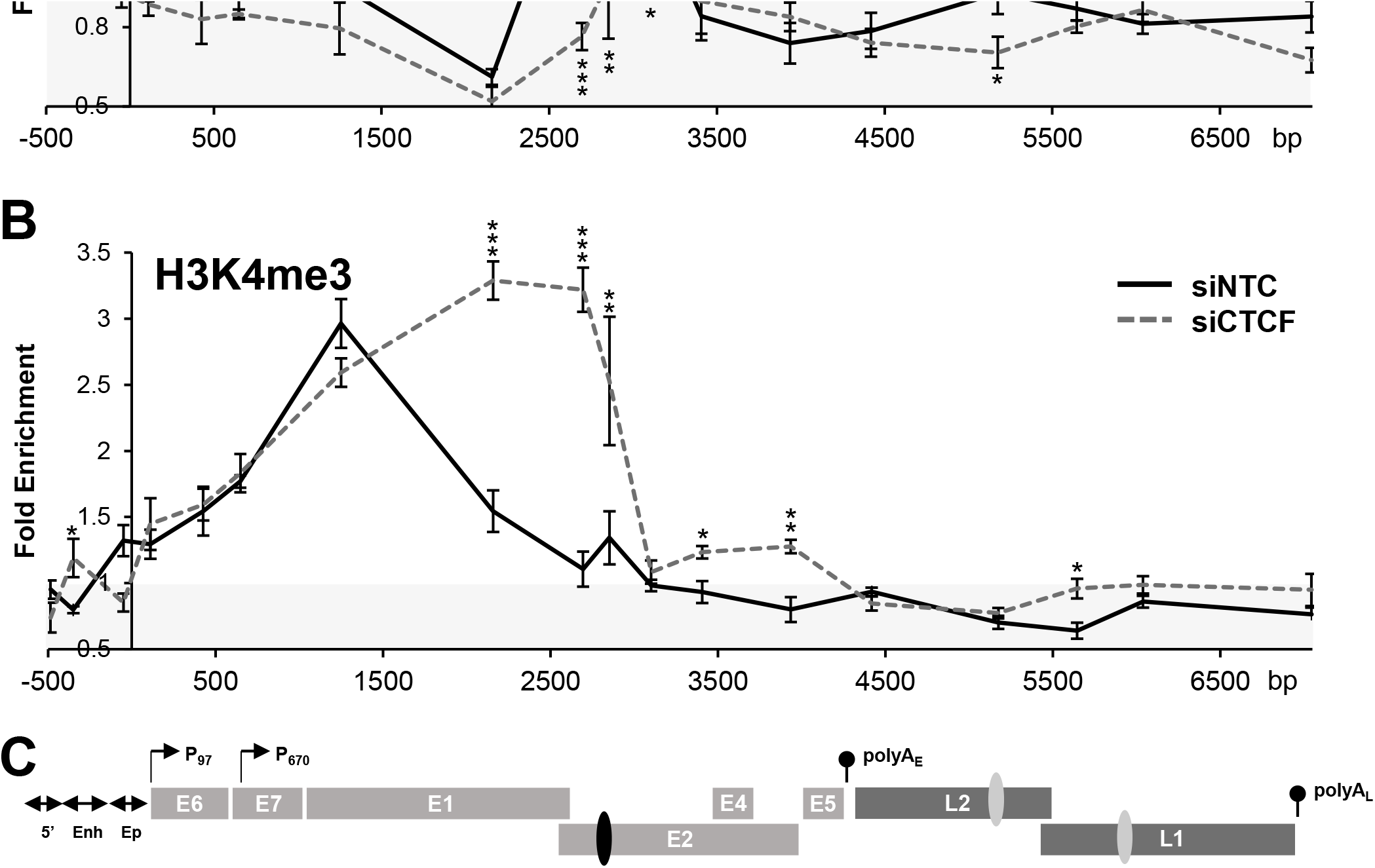
CTCF knockdown in W12 episomal cells is associated with elongated transcriptional activity across the HPV16 genome. Chromatin immunoprecipitation analysis of (A) CTCF and (B) active transcriptional mark H3K4me3 across (C) the episomal HPV16 genome from undifferentiated monolayer W12 episomal cells treated with non-targeting control (NTC, black lines) or siRNA specific to CTCF (dark grey hatched lines). In each graph, the y-axis shows the relative levels of enrichment above IgG controls (<1, grey area). Data points = mean±s.e.m. of n=3. Student’s T-test vs siNTC data: *p<0.05, **p<0.01, ***p<0.001.

**Figure 6.**
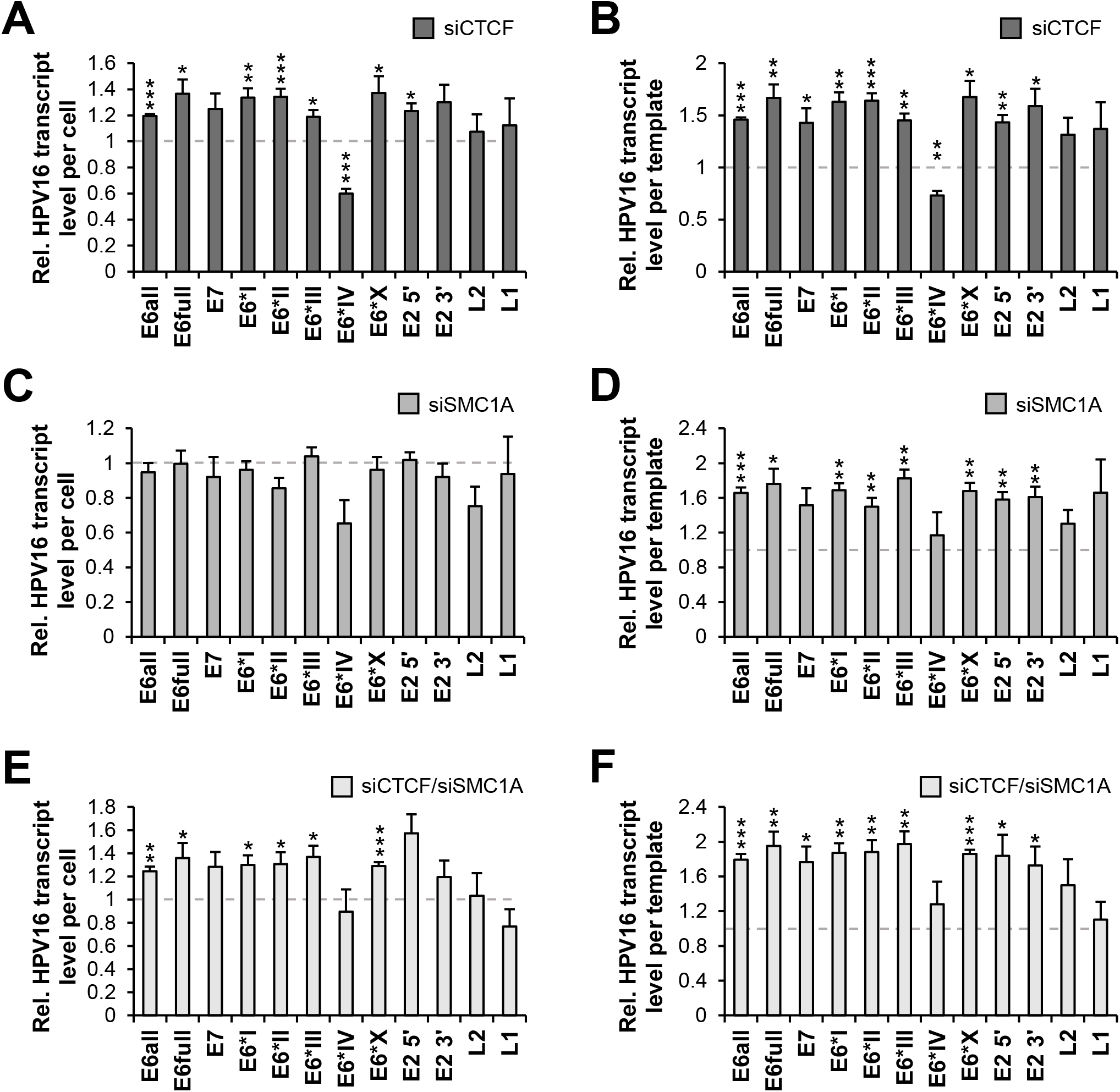
CTCF and SMC1A knockdown affect HPV16 transcript level in undifferentiated monolayer W12 episomal cells. Monolayer W12 episomal cells were treated with siRNAs specific to (A, B) CTCF (dark grey bars), (C, D) SMC1A (medium grey bars) and (E, F) CTCF/SMC1A (light grey bars) with HPV16 transcript level determined per cell (left column) and per virus template (right column), relative to non-targeting control (NTC) treated cells. Bars = mean+s.e.m. of n=3. Student’s T-test vs siNTC data: *p<0.05, **p<0.01, ***p<0.001.

## Discussion

Although the role of CTCF association with the episomal HPV18 genome has been well characterised, the function of CTCF interaction with the episomal HPV16 genome remains less well understood. Here, we show that, during methylcellulose-mediated differentiation of naturally infected W12 episomal cells, previously defined enrichment of CTCF at the HPV16 *E2* CTCFbs increases concurrently with modulation of virus early transcript levels and induction of late gene expression. Consistent with epithelial differentiation, HPV16 genome copy number per cell increases as expected with genome amplification during the virus life cycle. Indeed, when taken into account, data clearly demonstrate an overall early gene transcript decrease after 48 hours differentiation whilst late gene transcripts are still increasing, concordant with upper layers of HPV infected differentiating epithelium. Heightened transcript levels across the course of differentiation are also mirrored by increased association of the histone PTM H3K4me3 beyond the HPV16 late promoter, consistent with activation of transcription from this promoter.

The incremental enrichment of CTCF at the HPV16 *E2* CTCFbs here does contrast with the loss of CTCF association at the HPV18 genome during differentiation seen elsewhere[16]. However, different profiles of CTCF association with other HPV genomes (e.g. HPV31) genomes have previously been observed[15] and the methodology of methylcellulose differentiation used here is distinct from the transfection of keratinocytes with HPV genomes and organotypic raft culture or calcium-mediated differentiation of HPV infected cells used elsewhere[13, 15, 16]. Hence, technical differences could result in differing data profiles. However, our transcript data is consistent with expected patterns of HPV gene expression during epithelial differentiation[6, 10]. Moreover, the increasing association of CTCF at the *E2* CTCFbs during differentiation correlates directly with the sustained increase in E6*IV transcript level. Interestingly, HPV16 E6*IV is the only virus transcript that encodes E6 and full length E2 protein without E7, and is dependent on a splice acceptor site (2708 nt) in close proximity to the *E2* CTCFbs (2915 nt). Importantly, the ablation of CTCF in monolayer, undifferentiated W12 cells caused the converse effect to E6*IV transcript level, providing evidence that the presence of CTCF at the *E2* CTCFbs is essential for E6*IV expression. This finding is consistent with CTCF binding at the *E2* CTCFbs in HPV18 genomes, which regulates the level of the HPV18 233^3434 E6*II spliced transcript[13] and may be due to the ability of CTCF to cause slowing or pausing of RNA polymerase II (RNAPII) progression, facilitating inclusion of upstream exons as has been shown with the cellular *CD45* gene[32]. As E7 protein increases genomic instability[33], E6*IV expression may be a mechanism by which HPV16 avoids excessive host genome instability and possible negative effects on virus life cycle. Another protein translated from the E6*IV transcript is E5, which is known to have many functions, including genome amplification through its actions on the EGFR signalling pathway[6, 34]. Since HPV18 E6*II and HPV16 E6*IV both encode E5, it is possible that CTCF may regulate E5 protein levels in episomal cells such that increasing E5 protein synthesis confers an advantage for virus episome maintenance and replication, although without virus protein expression level analysis here this remains untested.

The incremental recruitment of CTCF to HPV16 genomes during methylcellulose-mediated differentiation of W12 cells also occurred at the putative *L2* CTCFbs and coincided with induction of late gene expression. Here, late gene expression was assessed by analysis of total L2 and L1 transcripts driven from the HPV16 late promoter, but also the spliced E4^L1 transcript which excludes the *L2* exon. As with recruitment at the *E2* CTCFbs, CTCF binding to the *L2* CTCFbs could result in upstream exon inclusion, i.e. the *E4* exon, or coincident binding at *E2* and *L2* CTCFbs could lead to loop formation, which may aid splicing between the *E4* and *L1* genes and lead to the >100-fold increase in this transcript seen with cellular differentiation, as well as allowing the induction of full length L2 and L1 transcripts.

Notably, SCM1A association at the *L2* CTCFbs remained unchanged during differentiation, similar to work performed elsewhere with HPV31[15]. Co-association of CTCF and SMC1A at the *L2* CTCFbs site of HPV31 is well characterised and supports virus genome maintenance in monolayer cells and amplification in differentiating cells[15]. Indeed, our data support the need for both components at HPV genomes in undifferentiated cells, as ablation of CTCF and SMC1A individually led to decreases in HPV16 genome copy number after 48 hour treatment, with dual ablation giving an intermediate, rather than additive, effect. Although necessity for these factors during differentiation-associated genome amplification was not examined here, it is clear across studies that recruitment of these factors throughout the HPV life cycle is critical for orderly control of virus genome replication[15]. Loss of episomal HPV genomes could occur through integration into the host genome[15], though this should not immediately result in a decreased copy number, which occurs with selection of HPV integrants[35]. Interestingly, a correlation between HPV16 integration sites and CTCF binding sites in the host genome has been established[31], and interaction between SMC1A and ChlR1, a known tethering factor of HPV genomes is known to occur[36, 37]; whether this association of CTCF and integration sites occurs due to previous tethering here is unknown. Furthermore, disruption of SMC1A and/or CTCF in HPV31 carrying CIN-612 cells and subsequent loss of genomes has also been hypothesised to be related to disruption of HPV infection mediated activation of the ataxia telangiectasia mutated-dependent (ATM) DNA damage response and the ataxia telangiectasia mutated-dependent DNA-related (ATR) pathway[15, 38, 39]. Here, SMC1A recruits DNA damage binding proteins to replicating virus genomes to maintain orderly replication, which could be dependent on initial CTCF binding of HPV sequences. Hence, these factors are likely to play a hierarchical, broader role during the HPV life cycle.

Indeed, the importance of CTCF during regulation of HPV transcription is clearly visible here. Increased CTCF association across the HPV16 genome during cellular differentiation and ablation of CTCF interaction in monolayer, undifferentiated W12 cells both cause initial generalised increases in virus early gene expression. Thus, it is likely that CTCF is acting in at least two different manners, besides upstream exon inclusion, while bound to the HPV16 genome. CTCF is known to insulate regions of active transcription from repressed regions of chromatin across the human genome[14]. Therefore, ablation of CTCF interaction with the *E2* CTCFbs in monolayer W12 cells may lead to early promoter activity upregulation, consistent with an increase in the transcriptional active histone PTM H3K4me3 across the early genes. CTCF also insulates cellular promoters from their enhancers, at least in part through causing genome loops[14]; a process that may cause aberrant cellular gene expression changes through HPV16 genome integration and insertion of a virus CTCFbs[31, 35]. This looping phenomenon has also been shown within HPV18 genomes between CTCF and YY1 to regulate the virus early transcript level, before differentiation of cells release some repression[16]. Hence, it is not surprising that disruption of CTCF binding to the HPV16 genome in monolayer W12 cells results in an increase in virus early transcript level here. Importantly, late gene expression does not appear to be activated, which is consistent with the necessity of cellular differentiation and multi-layered epigenetic changes to the HPV genome to additionally drive late promoter activation[40]. The finding that SMC1A ablation only statistically significantly increases virus early transcript levels with either co-ablation of CTCF or when HPV16 genome copy number is taken into account, underlines that CTCF appears to be the dominant player in control of HPV16 gene expression.

Taken together, our study shows that, whilst SMC1A association with the episomal HPV16 genome is important for genome maintenance and amplification, interaction of CTCF additionally is integral for regulation of virus transcription and splicing throughout the HPV life cycle. This control likely safeguards host cells from an excessive level of virus oncoproteins E6 and E7, which could more readily lead to tumourigenesis and a loss of full productive infection for the virus itself. Moreover, CTCF association with HPV genomes is likely to play a role in the reduced risk of cancer development during episomal HPV16 infections as opposed to those associated with integrated HPV genomes.

## Acknowledgements

Research was funded by a Cancer Research UK Programme Award (A13080).

## Author contributions

IJG and NC conceived the study. IJG devised the experimental approach. IJG, GT and IP performed the experiments. IJG, GT and IP analysed the data. IJG wrote the manuscript with GT, JLP and NC editing the manuscript.

